# Distinct neuroinflammatory signatures exist across genetic and sporadic ALS cohorts

**DOI:** 10.1101/2023.01.19.524561

**Authors:** Olivia M. Rifai, Judi O’Shaughnessy, Owen R. Dando, Alison F. Munro, Michael D.E. Sewell, Sharon Abrahams, Fergal M. Waldron, Christopher R. Sibley, Jenna M. Gregory

## Abstract

Amyotrophic lateral sclerosis (ALS) is a neurodegenerative disease characterised by progressive loss of upper and lower motor neurons. ALS is on a pathogenetic disease spectrum with frontotemporal dementia (FTD), with patients sometimes experiencing elements of both conditions (ALS-FTSD). For mutations associated with ALS-FTSD, such as the *C9orf72* hexanucleotide repeat expansion (HRE), the factors influencing where an individual may lie on this spectrum require further characterisation. Here, using NanoString molecular barcoding with a panel of 770 neuroinflammatory genes, we interrogate inflammatory dysregulation at the level of gene expression. We identified 20 dysregulated neuroinflammatory genes in the motor cortex of deeply clinically phenotyped C9-ALS *post-mortem* cases, with enrichment of microglial and inflammatory response gene sets. Our analyses also revealed two distinct ALS-related neuroinflammatory panel signatures (NPS), NPS1 and NPS2, delineated by the direction of expression of proinflammatory, axonal transport and synaptic signalling pathways. Two genes with significant correlations to available clinical metrics were selected for validation: *FKBP5* and *BDNF*. FKBP5 and its signalling partner, NF-*κ*B, appeared to have a cell-type-specific staining distribution, with activated (i.e., nuclear) NF-*κ*B immunoreactivity in C9-ALS. Expression of *BDNF*, a correlate of disease duration, was confirmed to be higher in individuals with long compared to short disease duration using BaseScope™ *in situ* hybridisation. Finally, we compared NPS between C9-ALS cases and those from deeply clinically phenotyped sporadic ALS (sALS) and SOD1-ALS cohorts, with NPS1 and NPS2 appearing across all cohorts. A subset of these signatures was also detected in publicly available RNA-sequencing data from independent C9-ALS and sALS cohorts, underscoring the relevance of these pathways across cohorts. Our findings highlight the importance of tailoring therapeutic approaches based on distinct molecular signatures that exist between and within genetic and sporadic cohorts.

## Introduction

Hexanucleotide repeat expansions (HREs) in the *C9orf72* gene are one of the most common mutations associated with amyotrophic lateral sclerosis (ALS) and ALS-frontotemporal spectrum disorder (ALS-FTSD)^1–4^. Clinical manifestations of disease associated with *C9orf72* HRE are variable; presentations can involve motor or cognitive symptoms related to ALS-FTSD, or other symptoms such as parkinsonism and psychosis^5–8^. This heterogeneity occurs despite a seemingly unifying neuropathological phenotype characterised by p62, TDP-43 and dipeptide repeat protein (DPR) deposits^9–14^. Clinical heterogeneity has the potential to be a large confounding factor in clinical trials including people with ALS-FTD, which often employ outcome measures based on clinical phenotypes. Thus, a better understanding of the molecular mechanisms underlying the clinical heterogeneity seen in C9-ALS, and people with ALS-FTD generally, is critical for informing the design of therapeutics intended to reduce specific symptom burden, as well as for improved trial stratification, so that endpoints can be more meaningfully measured.

One potential factor influencing variable disease presentation in people with ALS-FTD is immune function and its related inflammatory processes. Inflammatory factors such as regulatory T cells and interleukins have previously been shown to be associated with the rate of disease progression^15–17^. Furthermore, differences in neuroinflammatory markers like CHIT1 and GFAP have been observed in the cerebrospinal fluid between ALS, FTD and ALS-FTSD patients^18,19^, suggesting that differential processes, particularly those regulated by neuroglia, may be occurring between conditions^18^. As *C9orf72* is highly expressed in microglia^20^, the resident immune cells of the central nervous system, it has been suggested that microglia may be particularly susceptible to any negative consequences of a change in normal C9orf72 protein function, thus triggering immune dysfunction^21^, as evidenced by knockout *C9orf72* models^22,23^. We have previously shown with immunohistochemical staining of *post-mortem* tissue that microglial activation is elevated in the language-related region Brodmann area (BA) 39 in language-impaired C9-ALS cases^24^. Additionally, we have demonstrated with random forest modelling that microglial staining is an accurate classifier of C9-ALS, with better sensitivity and specificity to disease than other markers such as astrocyte activation marker, GFAP, and phosphorylated TDP-43 aggregate marker, pTDP43^24^. Thus, further characterisation of immune dysfunction and its influence on clinical heterogeneity in C9-ALS, especially at a molecular level, is warranted to understand how these pathways can be more specifically targeted to harness their therapeutic potential.

To date, few studies have taken a targeted approach to measuring the expression of neuroinflammatory genes in a C9-ALS cohort, particularly in *post-mortem* tissue. One recent study observed a general enriched immune response in *post-mortem* frontal cortex tissue from *C9orf72* HRE carriers^25^, though this response was not explored further as the focus of the study. To interrogate inflammatory dysregulation in this context at a molecular level, we performed NanoString molecular barcoding on *post-mortem* motor cortex from a cohort of deeply clinically phenotyped C9-ALS and C9-ALS-FTSD cases to explore differential expression of 770 neuroinflammatory genes. We identified 20 significantly differentially expressed neuroinflammatory genes in a deeply clinically phenotyped C9-ALS *post-mortem* tissue cohort, revealing clustering of therapeutically relevant gene expression patterns. We compared gene expression patterns with immunohistochemical data from our previous study to examine relationships between gene dysregulation and neuropathological staining^24^ We also performed regional validation of two genes correlating with clinical scores using both immunohistochemical and BaseScope™ *in situ* hybridisation techniques. Finally, we identified similarities and differences in inflammatory signatures between C9-ALS, sporadic ALS (sALS), and SOD1-ALS cohorts, as well as in publicly available frontal cortex RNA sequencing data from independent C9-ALS and sALS cohorts^26^.

## Methods

### Case identification and cognitive profiling

*Post-mortem* tissue from cases with ALS or ALS-FTSD (*n = 33*) was obtained from the Medical Research Council (MRC) Edinburgh Brain Bank (Table 1, Table 3). For genetic classification of all ALS cases, repeat-primed polymerase chain reaction (PCR) was carried out for *C9orf72* HRE identification and whole genome sequencing was carried out to identify other ALS-associated mutations. SOD1-ALS cases were confirmed to have an I114T missense mutation^27^. Sporadic cases had no family history of ALS, and no ALS-associated mutations identified through gene panel analysis^27^. *Post-mortem* tissue from controls age- and sex-matched to C9-ALS cases (*n = 10*) with no history of neurological conditions or neurodegenerative pathology was obtained from the Edinburgh Sudden Death Brain Bank. *Post-mortem* tissue was collected with ethics approval from East of Scotland Research Ethics Service (16/ES/0084) in line with the Human Tissue (Scotland) Act (2006); the use of *post-mortem* tissue for studies was approved by the Edinburgh Brain Bank ethics committee and the Academic and Clinical Central Office for Research and Development (ACCORD) medical research ethics committee (AMREC). Clinical data were collected for the Scottish Motor Neurone Disease Register (SMNDR) and Care Audit Research and Evaluation for Motor Neurone Disease (CARE-MND) platform, with ethics approval from Scotland A Research Ethics Committee (10/MRE00/78 and 15/SS/0216). Donor patients underwent neuropsychological testing with the Edinburgh Cognitive and Behavioural ALS Screen (ECAS)^28^. Clinical correlates of motor dysfunction/disease progression include disease duration (months) and sequential ALS functional rating scale (ALSFRS) data points. Clinical correlates of cognition include Edinburgh Cognitive and Behavioural ALS Screen (ECAS) scores for ALS-specific and ALS non-specific subdomain scores. All patients consented to use of their data during life.

**Table 1.** C9-ALS cohort demographics.

| Case Number | Sex | Age at death (y) | Clinical Diagnoses | Disease duration (m) | Region of onset | ECAS | ALSFRS | Neuronal pTDP43 burden | Glial pTDP43 burden |
| --- | --- | --- | --- | --- | --- | --- | --- | --- | --- |
| C9-ALS 1 | F | 63 | ALS | 25 | Lower Limb | Yes (unimpaired) | No | 2 | 2 |
| C9-ALS 2 | M | 50 | ALS | 29 | Bulbar | Yes (language dysfunction) | Yes (4 data points) | 2 | 3 |
| C9-ALS 3 | F | 63 | ALS | 30 | Upper limb | No | Yes (3 data points) | 2 | 2 |
| C9-ALS 4 | F | 62 | ALS | 37 | Mixed | Yes (unimpaired) | Yes (6 data points) | 1 | 2 |
| C9-ALS 5 | F | 65 | ALS | 44 | Lower limb | No | Yes (2 data points) | 1 | 2 |
| C9-ALS 6 | M | 65 | ALS | 50 | Upper limb | No | Yes (1 data point) | 1 | 1 |
| C9-ALS 7 | M | 62 | ALS | 50 | Lower limb | Yes (executive dysfunction) | Yes (2 data points) | 2 | 0 |
| C9-ALS 8 | M | 43 | ALS | 57 | Bulbar | Yes (language dysfunction) | No | 1 | 2 |
| C9-ALS 9 | M | 58 | ALS | 87 | Lower limb | Yes (language dysfunction) | Yes (3 data points) | 3 | 3 |
| C9-ALS 10 | F | 63 | ALS, FTD, Schizophrenia | 119 | Upper limb | Yes (FTD – 3 domains) | No | 2 | 3 |
| Control 1 | F | 65 | Hypertension, Cardiomyopathy | N/A | N/A | N/A | N/A |  |  |
| Control 2 | M | 50 | None | N/A | N/A | N/A | N/A | 1 | 0 |
| Control 3 | F | 57 | None | N/A | N/A | N/A | N/A | 0 | 0 |
| Control 4 | F | 57 | None | N/A | N/A | N/A | N/A | 0 | 0 |
| Control 5 | F | 71 | Depression, Paranoid Schizophrenia | N/A | N/A | N/A | N/A | 0 | 0 |
| Control 6 | M | 58 | Depression | N/A | N/A | N/A | N/A | 1 | 0 |
| Control 7 | F | 59 | Hypertension | N/A | N/A | N/A | N/A | 0 | 0 |
| Control 8 | M | 44 | None | N/A | N/A | N/A | N/A | 0 | 0 |
| Control 9 | M | 63 | None | N/A | N/A | N/A | N/A | 0 | 0 |
| Control 10 | F | 61 | Depression, MS, hypothyroidism | N/A | N/A | N/A | N/A | 0 | 0 |

### NanoString sequencing and analysis

RNA from human tissue was extracted using the RNAstorm FFPE RNA extraction kit (Cell Data Sciences, Fremont, CA, USA) on two 10 µm curls per sample cut from BA4. RNA was eluted in 50 uL nuclease-free water, after which sample concentrations were measured using a NanoDrop 1000 (ThermoFisher Scientific, Waltham, MA, USA). Samples that did not meet the minimum 60 ng/uL were concentrated using an Eppendorf Concentrator Plus (Eppendorf, Hamburg, Germany) for 10 minutes at 45°C, measured again, and concentrated for an additional 5 minutes at 45°C if necessary. Samples were diluted in nuclease-free water to a final concentration of 600 ng RNA in 10 uL water for NanoString sequencing. Sequencing was performed by Host and Tumour Profiling Unit (HTPU) Microarray Services with the nCounter neuroinflammation panel (for more information see https://nanostring.com/products/ncounter-assays-panels/neuroscience/neuroinflammation/), which includes 770 genes related to immunity and inflammation, neurobiology and neuropathology, and metabolism and stress^29^ (Supplementary Material). All samples passed routine quality control checks. Differential gene expression analyses between control and ALS cohorts were performed in RStudio (R version 4.1.1)^30^ using “DESeq2”^31^ (version 1.32.0) with “RUVSeq”^32^ (version 1.26.0) to estimate and regress out unwanted variation (*k = 3* factors of unwanted variance^33^). P-values were adjusted using Benjamini-Hochberg false discovery rate thresholds (*p < 0*.*05*). Plots were made using the “ggplot2” package^34^ (version 3.3.5) in R. Gene ontology (GO) enrichment analysis^35^ was performed with the “topGO” package (version 2.44.0)^36^ in R, and gene set analysis correlation adjusted mean rank (CAMERA)^37^ from the “limma” package^38^ (version 3.48.3) in R was performed using the Molecular Signatures Database (MSigDb)^39,40^ GO category terms, with the nCounter neuroinflammation gene panel as the background gene set and gene annotations taken from Ensembl (version 96)^41^. GO and CAMERA results were considered significant if -log10(p-value) > 1.3; significant CAMERA results were filtered for gene sets with *n = 10*+ genes. Clustering analyses were performed using the “pheatmap” package^42^ (version 1.0.12) in R. Correlations of IHC and ECAS data with housekeeping-normalised counts were calculated with the “corrplot” package^43^ (version 0.91) in R, using Spearman’s test. Gene set testing was also performed with cell-type-specific gene sets derived from published Brain RNA-seq data^20^ to determine cell-type-specific dysregulation of transcripts. For this analysis, ratios were calculated for the expression of each gene in each cell type compared to its maximum expression in any other cell type. ‘Human_(cell type)_5_times’ indicates all genes for which the calculated ratio is >5, ‘human_(cell type)_10_times’ indicates all genes for which the calculated ratio is >10, and ‘human_(cell type)_top100’ indicates the 100 genes with the highest ratio for that cell type, that is, the most specific genes for each cell type. Differential expression analysis between NPS1 and NPS2 cases in the C9-ALS cohort was conducted as above, and all genes with an unadjusted p-value < 0.05 and adjusted p-value ≠ NA were taken through clustering analyses with the “pheatmap” package^42^ (version 1.0.12) in R.

### Public RNA sequencing data analysis

A raw count matrix of publicly available frontal cortex and cerebellum RNA sequencing data^26^ were accessed via the NCBI Gene Expression Omnibus (accession number GSE67196). Data were divided by brain region and counts for C9-ALS and sALS cases were extracted. Data were variance stabilised and scaled (i.e., z-transformed) across samples using “DESeq2”^31^ (version 1.32.0) in RStudio (R version 4.1.1)^30^. Clustering analyses were performed for each region using the “pheatmap” package^42^ (version 1.0.12) in R. Heatmaps included equivalent demographic, clinical or pathological information available with the public data analysed: sex, region of onset, and disease duration.

### Immunohistochemistry (IHC)

*Post-mortem* brain tissue was obtained from Brodmann areas (BA) BA4, BA39, BA44, BA46 and fixed in 10% formalin for a minimum of 72 h. These regions were selected for their associations with clinical phenotypic correlates as we have shown previously^28^; BA4 – motor, BA39 – language, BA44 – fluency and language, BA46 – executive function. For this validation dataset, an additional case and alternate control (Case 11 and Control 11 in Supplementary Materials) were included due to differences in tissue availability at the time of request. Tissue was dehydrated in a 70–100% ascending alcohol series and subsequently washed three times for 4 hours in xylene. Three 5-hour paraffin wax embedding stages were performed, after which formalin-fixed, paraffin-embedded (FFPE) tissue was cooled and sectioned on a microtome (ThermoFisher Scientific) into 4 μm serial sections. Sections were placed on Superfrost (ThermoFisher Scientific) microscope slides and left to dry overnight at 40°C. Sections were dewaxed with successive xylene washes, hydrated with alcohol, and treated with picric acid to remove formalin pigment and quench lipofuscin. For NF-*κ*B staining, antigen retrieval was carried out in Tris-EDTA buffer (pH 9) in a pressure cooker for 30 min, after which a Novolink Polymer detection system^44^ was used with an Abcam anti-NF-*κ*B antibody (Abcam, Cambridge, UK) at a 1 in 1500 dilution. For FKBP5 staining, antigen retrieval was carried out in citric acid buffer (pH 6) in a Pressure King Pro pressure cooker for a 20 min cycle; samples were heated to 140°C and incubated for 5 min, after which pressure was manually released. The Novolink Polymer detection system (Leica Biosystems, Newcastle, UK) was then used with an anti-FKBP5 antibody (OriGene, Rockville, MD, USA) at a 1 in 80 dilution. Staining was performed with 3,3’- diaminobenzidine (DAB) chromogen and counterstained with haematoxylin, as per standard operating procedures, after which slides were dehydrated, washed in xylene, and coverslips mounted using DPX mountant (Sigma Aldrich, St. Louis, MO, USA). For sequential staining, slides initially stained with NF-*κ*B or FKBP5 were soaked in xylene overnight, after which the coverslips were carefully removed, and the slides were soaked for several more hours until the DPX mountant had dissolved off the sections. Slides were re-stained according to standard operating procedures mentioned above, from hydration, to citric acid antigen retrieval with a pressure cooker, to staining with an anti-Iba1 antibody (Abcam) at a 1 in 3000 dilution. Protocols for CD68, Iba1, pTDP43 and GFAP staining were described previously^24^. Manual grading of neuronal and glial TDP-43 burden was performed by a pathologist (JMG) using a scale from 0 to 3 as outlined in a previous study^45^.

### Image analysis

For analysis of NF-*κ*B and FKBP5 immunohistochemical staining, whole tissue sections were scanned with Brightfield at 40x magnification using a Hamamtsu NanoZoomer XR (Hamamatsu Photonics (UK) Ltd, Welwyn Garden City, UK). Using NDP.view2 viewing software (Hamamatsu), regions of interest (ROIs) were taken from key regions for quantification. Three ROIs were taken from grey matter regions including layer V neurons, and three ROIs were taken from white matter regions. ROIs were analysed with QuPath software^46^ cell segmentation; cells were segmented using a watershed method based on haematoxylin counterstaining, with different parameters for grey and white matter and for neurons and glia to best distinguish between cell types. Full scripts used for the automated cell segmentation and quantification of NF-*κ*B and FKBP5 are included in Supplementary Information. Cells were classified as nuclear- and/or cytoplasmic-positive for each stain based on the DAB mean intensity of each compartment. Measurements were exported at the image (number of nuclear- and/or cytoplasmic-positive cells) and cell level (intensity and morphological features). Data were visualised in RStudio with the “ggplot2” package^34^ (version 3.3.5). Data were found to be non-normal via Shapiro-Wilk’s test and subjected to non-parametric tests (i.e., Mann-Whitney *U* for two-group comparisons, pairwise Wilcoxon for three-group comparisons and Spearman’s for correlations). Comparisons were only conducted between groups with *n ≥ 3*. Results were presented as ungrouped or grouped by brain region, grey or white matter, and vascular or non-vascular adjacent. Analysis methods for CD68, Iba1, pTDP43 and GFAP staining can be found in our previous study^24^.

### BaseScope^™^ in situ hybridisation

*In situ* hybridisation was performed on tissue sections using BaseScope reagents (Advanced Cell Diagnostics, Newark, CA, USA) as per the manufacturer’s instructions^47^ and as described previously^48^. Probe hybridisation was performed using BaseScope^™^ probes for *BDNF* mRNA transcripts. Slides were counterstained using haematoxylin and lithium carbonate, washed in xylene, and coverslips were mounted using DPX Mountant. For BDNF BaseScope^™^ *in situ* hybridisation, due to the sparsity of mRNA transcripts, expression was quantified manually by a pathologist (JMG), who was blinded to clinical data, and a second rater (OMR), who performed a 20% validation check, no discrepancies were identified between raters. Expression was quantified using a product score comprising of two factors: total transcript count per high-power field (40X) x total number of transcript-positive cells per high-power field. Ten high-power fields were evaluated and averaged across each case.

## Results

### Two distinct neuroinflammatory signatures exist in C9-ALS

To explore *C9orf72* mutation-related changes to glial activation at the gene expression level, motor cortex tissue was sequenced using NanoString molecular barcoding, which provides accurate mRNA counts without the need for amplification steps that favour highly abundant transcripts^29^. This method circumvents RNA degradation issues related to *post-mortem* autolysis, as probes bind to the central, most preserved portion of mRNA transcripts. A panel of 770 neuroinflammation-related genes was used, and differential expression analyses were conducted; raw and processed gene expression data are available in the Supplementary Information. The analysis revealed a list of 20 genes that were significantly differentially expressed between C9-ALS cases and controls (Figure 1a). The microgliosis we observed previously in C9-ALS^24^ is supported here by the upregulation of *CD163*, a marker of macrophage activity, and the downregulation of *P2RY12*, a marker of microglial homeostasis^49,50^. The 20 significantly differentially expressed genes clustered into two similarly sized groups, those that were upregulated in C9-ALS (*SERPINA3, S100A10, FKBP5, EMP1, CD163, SPP1, CP, CTSE, BAG3*), and those that were downregulated (*ARC, RALB, EGR1, JUN, COX5B, P2RY12, BDNF, SLC17A6, BAD, MFGE8, FOS*) relative to control cases. GO enrichment analysis revealed associations of these significantly dysregulated genes with pathways implicated in processes such as AP-1 complex signalling, pri-miRNA transcription, Smad-signalling, neuron projection and death, post-translational protein modification, and acute-phase response (Figure 1b). Importantly, as the number of significantly dysregulated genes in this analysis is relatively low, the GO findings must be interpreted with caution. Thus, we also employed a competitive gene set analysis, CAMERA, which considers whole shifts in expression of groups of genes based on fold changes. Neuron development, projection, and differentiation, gene sets were downregulated in C9-ALS cases relative to controls, as well as gene sets for synaptic structure, plasticity and transmission, cell projection organisation, and cytochrome C release; blood microparticle, platelet degranulation, endopeptidase inhibitor activity, and inflammatory response gene sets were upregulated (Figure 1c). Finally, microglia-specific genes were found to be significantly upregulated in our dataset, in line with our previous findings^24^, further supporting an increase in microglial activation^24^ (Table 2).

**Table 2.** Cell-type-specific dysregulation in C9-ALS based on Brain RNA-seq data.

| gene_set | NGenes | Direction | PValue | FDR |
| --- | --- | --- | --- | --- |
| human_microglia_5_times | 113 | Up | 8.58E-04 | 0.03089637 |
| human_microglia_10_times | 78 | Up | 0.00983159 | 0.08848435 |
| human_endothelial_top100 | 12 | Up | 0.02976191 | 0.13392859 |
| human_microglia_top100 | 38 | Up | 0.03978398 | 0.15913592 |
| human_endothelial_5_times | 18 | Up | 0.04770685 | 0.17174467 |
| human_endothelial_10_times | 13 | Up | 0.05512601 | 0.18041241 |
| human_fetal_astrocytes_5_times | 46 | Down | 0.13938532 | 0.34892127 |
| human_neuron_5_times | 19 | Down | 0.21064688 | 0.42102263 |
| human_fetal_astrocytes_10_times | 22 | Down | 0.22756045 | 0.42102263 |
| human_neuron_top100 | 5 | Down | 0.24559653 | 0.42102263 |
| human_neuron_10_times | 8 | Down | 0.27457446 | 0.44930366 |
| human_fetal_astrocytes_top100 | 7 | Down | 0.67226571 | 0.84000835 |
| human_mature_astrocytes_5_times | 9 | Up | 0.67667339 | 0.84000835 |
| human_oligodendrocyte_5_times | 24 | Down | 0.73334664 | 0.86196644 |
| human_mature_astrocytes_10_times | 5 | Up | 0.76421218 | 0.86196644 |
| human_mature_astrocytes_top100 | 7 | Down | 0.76619239 | 0.86196644 |
| human_oligodendrocyte_top100 | 21 | Up | 0.80590044 | 0.8624924 |
| human_oligodendrocyte_10_times | 14 | Up | 0.81457615 | 0.8624924 |

**Figure 1.**
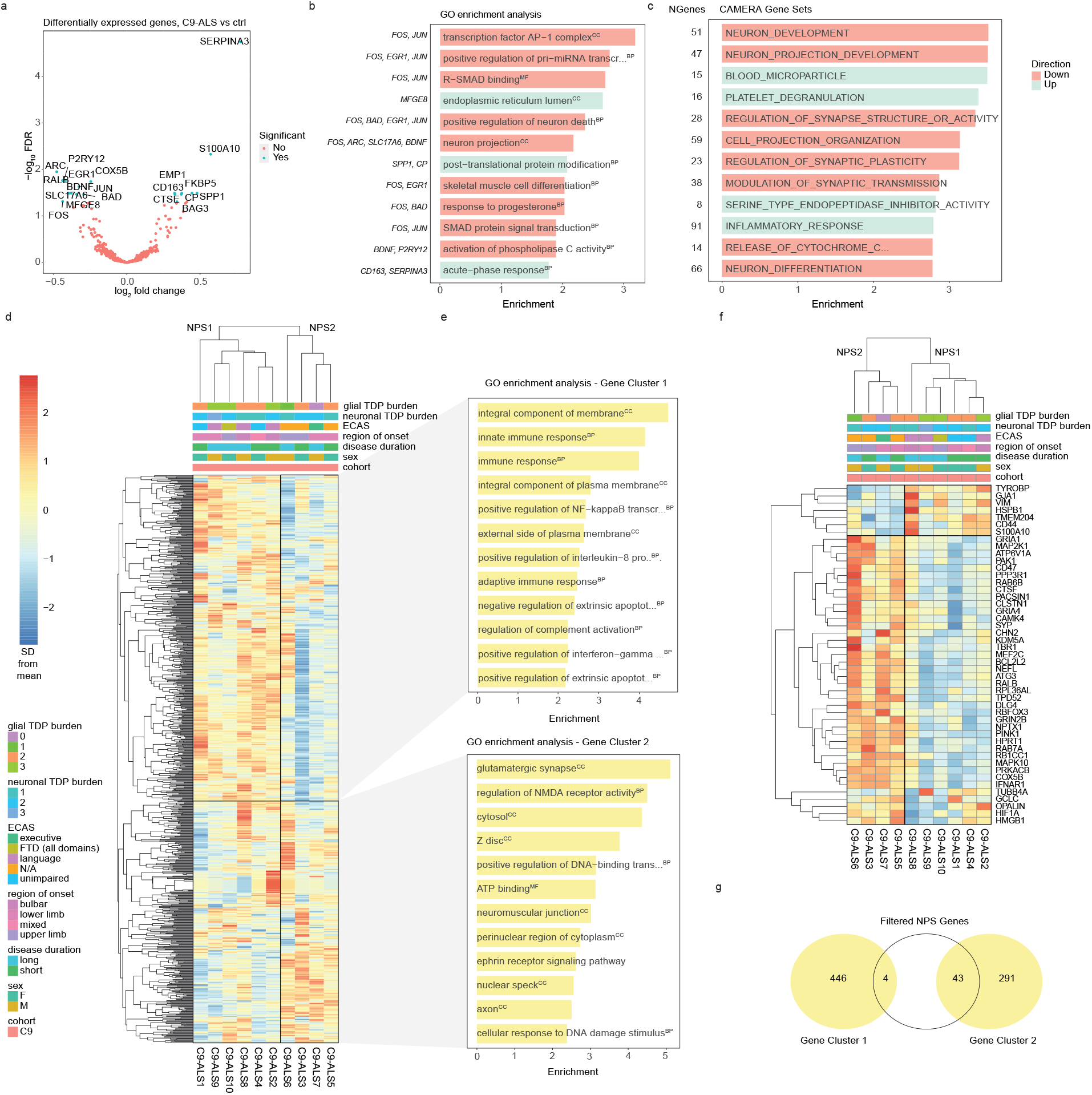
Two distinct neuroinflammatory signatures exist in C9-ALS. (a) Volcano plot showing differentially expressed genes between C9-ALS cases and controls by log2 fold change and -log10 p-value. Non-significant genes are represented in red and significant genes (i.e., surpassing the Benjamini-Hochberg false discovery rate (FDR) threshold of p-adjusted < 0.05) are represented in blue. (b) GO enrichment analysis of genes enriched in C9-ALS cases by type with -log10(p-value) score showing the top 12 most differentially expressed gene sets. Italicised terms indicate downregulation; key genes for each term are shown to the left; (c) CAMERA gene set analysis of gene sets dysregulated in ALS cases, with the number of genes for each set shown to the left, showing the top 10 most differentially expressed gene sets. (d) Clustered neuroinflammation panel heatmap including expression of entire NanoString panel (770 genes), showing two distinct neuroinflammatory panel signatures (NPS1 & NPS2) in C9-ALS. Gene clusters 1 & 2 are boxed, with opposite directions of expression between NPS. Clinical (region of onset, disease duration, ECAS) and pathological (neuronal and glial pTDP-43 burden) keys are shown. (e) GO enrichment analysis for Gene Cluster 1 and 2 (that define NPS1 and NPS2) showing top 12 most differentially expressed gene sets. Italicised terms indicate downregulation; (f) Clustered neuroinflammation panel heatmap including only differentially expressed genes between C9-ALS cases with NPS1 and NPS2, showing two distinct NPS, with genes listed on the right; (f) Venn diagram showing overlap between genes from the filtered NPS gene list in (e) with Gene Cluster 1 and 2 from (d-e). MF, molecular factor; CC, cellular component; BP, biological process.

Examination of gene expression differences in C9-ALS cases across the whole neuroinflammatory panel revealed the existence of two distinct gene expression signatures, herein referred to as neuroinflammatory panel signature 1 and 2 (NPS1 and NPS2); these signatures defined two disease clusters and were delineated by the direction of expression of two gene clusters (Figure 1d). The clearest phenotypic distinctions between NPS1 and NPS2 observed were that of manually graded glial TDP-43 burden and language impairment as determined by the Edinburgh Cognitive and Behavioural ALS Screen (ECAS), with highest TDP-43 burden and language impairment only occurring in NPS1. GO analysis of C9-ALS gene clusters revealed an enrichment of immune and inflammatory response pathways in gene cluster 1, such as positive regulation of interleukin-8, NF-*κ*B and interferon-gamma responses (Figure 1e). By contrast, gene cluster 2 exhibited an enrichment of axonal transport and synaptic signalling pathways (Figure 1d). To determine which genes within the panel were contributing to the delineation of these clusters, differential expression analysis was conducted to identify differentially expressed genes between cases exhibiting NPS1 and NPS2 expression signatures. Forty-seven genes were included in a new clustered heatmap (herein referred to as the NPS-defining gene list), exemplifying a clearer contrast between the direction of expression of genes between the two signatures (Figure 1f). These genes were mostly from original gene cluster 2, related to axonal transport and synaptic signalling (Figure 1g).

### Differentially expressed genes correlate with microglial and pTDP-43-related immunohistochemical features in C9-ALS

To explore the relationship of differentially expressed genes in C9-ALS with glial activation and TDP-43 burden (Figure 2a), we correlated microglia-, astrocyte-, and TDP-43-related immunohistochemical data from our previous digital pathology study^24^ with transcript counts of the 20 differentially expressed genes identified in Figure 1. These data consisted of digitally extracted features (i.e., stain-positive superpixel counts, a measurement of stain abundance) from stained motor cortex (BA4) of the same C9-ALS cohort included in the current study, and included Iba1 (i.e., homeostatic microglia), CD68 (i.e., activated macrophage), GFAP (i.e., activated astrocyte), and pTDP43 (i.e., phosphorylated TDP-43 aggregate) staining. Expression levels of several genes were found to correlate significantly with the number of CD68+ or pTDP43+ superpixels, with positive correlations between CD68+ and proinflammatory genes (e.g., FKBP5, CD163, SPP1), as well as with molecular chaperone regulator BAG3, and negative correlations between pTDP43 and the expression of JUN and FOS, subunits that form the transcription factor complex activator protein 1 (AP-1). (Figure 2b). When subdivided by disease status, the significant proinflammatory correlations with CD68+ were lost in controls, and a significant negative correlation of homeostatic microglia marker *P2RY12* expression with pTDP43+ appeared (Figure 2b). Finally, a positive correlation with expression of the growth factor *BDNF* with pTDP43+ superpixels was seen in C9-ALS but not controls (Figure 2b). Interestingly, when cases were divided by NPS, NPS1 cases exhibited more positive correlations with stain abundance, while NPS2 correlation coefficients were more often negative (though non-significant). These data suggest distinct NPS-related directionality in correlations between gene expression and pathological features such as TDP-43 aggregation and glial activation (Figure 2c), in line with our observation that C9-ALS cases with a predominance of NPS1 gene expression were more likely to have higher glial TDP-43 aggregation burden (Figure 1d).

**Figure 2.**
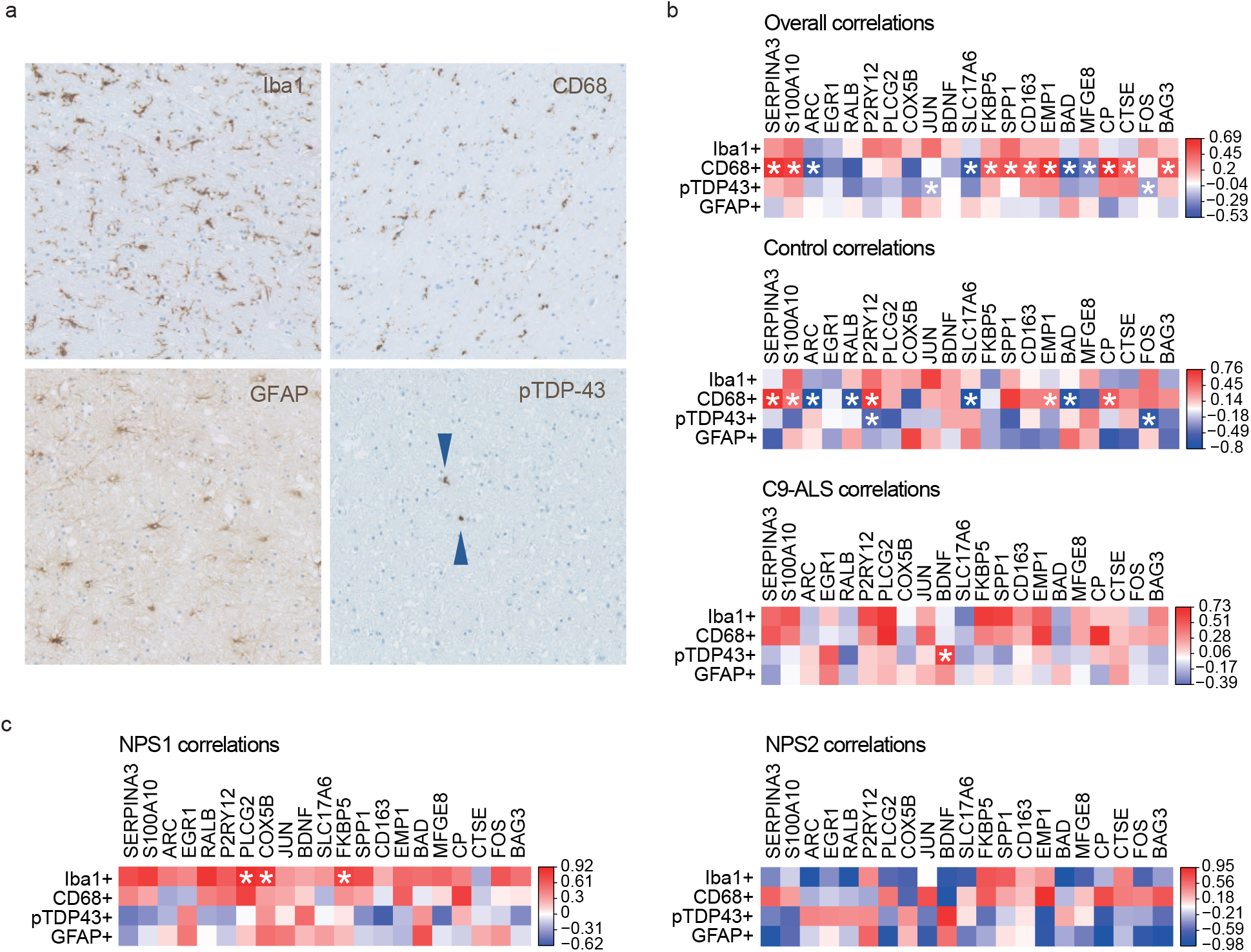
Differentially expressed genes correlate with microglial and pTDP-43-related immunohistochemical features in C9-ALS. (a) Example images of immunohistochemical stains quantified using QuPath for correlations with NanoString neuroinflammation panel housekeeping-normalised counts. (b) Control-specific, C9-ALS-specific, and overall significant correlations of prominence scores for Iba1, CD68, pTDP-43 and GFAP stains with NanoString normalised counts for 20 dysregulated genes in C9-ALS from Figure 1a. (d) NPS1- and NPS2-specific correlations of prominence scores for Iba1, CD68, pTDP-43 and GFAP stains with NanoString normalised counts for 20 dysregulated genes in C9-ALS from Figure 1a. Spearman’s R correlation coefficients are indicated by colour, and correlations with a p-value < 0.05 are marked with an asterisk.

### FKBP5 expression correlates significantly with clinical metric of executive dysfunction for C9-ALS

To investigate possible relationships between differential transcription patterns and cognition, we examined correlations between differential gene expression and ECAS scores, for the 20 genes we identified as most differentially expressed between C9-ALS cases and controls. (Figure 3a); *FKBP5* and *COX5B* were found to negatively correlate with executive score. These relationships must be interpreted with caution as gene expression was measured in the motor cortex and not regional correlates of ECAS scores; however, it may be that changes in the motor cortex are reflective of changes in the relevant regions and may reflect a global cortical burden of disease. The immunophilin FK506-binding protein 51 (FKBP5) modulates inflammation through nuclear factor- κB (NF-κB) signalling^51,52^, and forms a chaperone complex with a heat shock protein (HSP90) in response to stress^53^. We interrogated whether this was also the case in C9-ALS brain tissue by using immunohistochemistry, rather than *in situ* hybidisation, as the functional form of NF-κB is a protein whose cellular localisation and expression level determines its function. Serial tissue sections were stained with FKBP5 and NF-κB and compared between control and C9-ALS tissue. No evidence of a significant increase in nuclear/cytoplasmic FKBP5 intensity ratios was observed, though there was a general trend towards an increase in C9-ALS (Supplementary Figure 1). However, significant increases in nuclear/cytoplasmic NF-κB intensity ratios were found in BA4 grey matter in C9-ALS, suggesting upregulation of this pathway in disease (Figure 3b). Upon sequential staining of the same FKBP5- or NF-κB-stained tissue with Iba1, cell-type-specific staining was observed for both FKBP5 and NF-κB (Figure 3c). Notably, microglia were found to be FKBP5+, accompanied by both FKBP5+ and FKBP5- neuroglia of other subtypes. Contrastingly, microglia were the only glial subtype found to be NF-κB+ (Figure 3d). Finally, when cases were stratified by inflammatory signature, NPS1 cases exhibited significantly higher nuclear/cytoplasmic NF-κB ratios in grey matter glia in BA4, and significantly lower ratios in neurons in extramotor BA44 (Figure 3e).

**Figure 3.**
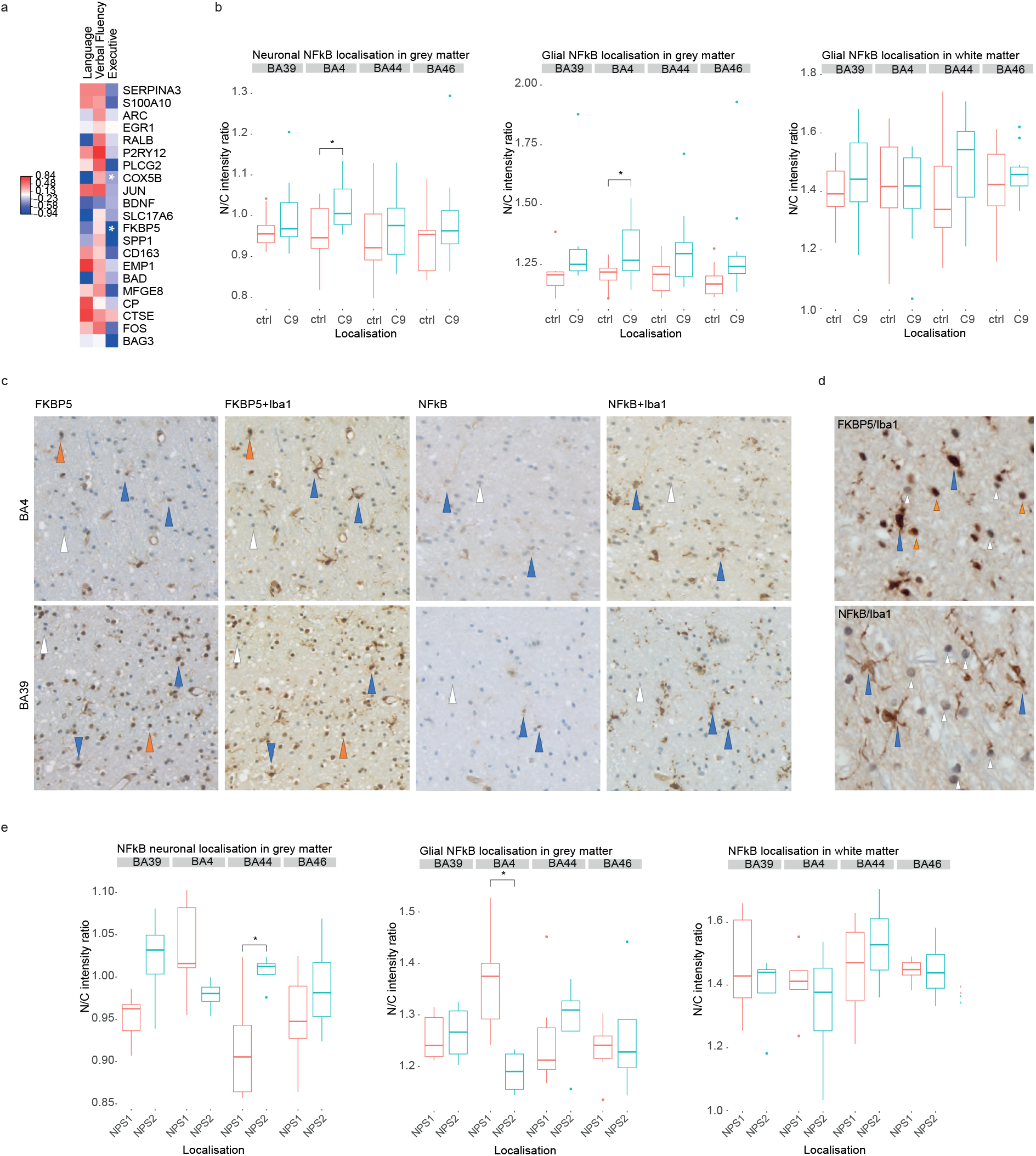
FKBP5 expression correlates significantly with clinical metric of executive dysfunction. (a) Correlation between language, fluency, and executive scores from ECAS and NanoString normalised counts for 20 dysregulated genes in C9-ALS from Figure 1a. Spearman’s R correlation coefficients are indicated by colour, and correlations with a p-value < 0.05 are marked with an asterisk. (b) Nuclear/cytoplasmic NF-*κ*B intensity ratio quantification for neuronal (left), grey matter glial (middle) and white matter glial (right) staining between ALS and controls, stratified by brain region. * *p < 0*.*05*. (c) Double staining with FKBP5 or NF-*κ*B and Iba1 to identify microglia-specific staining in BA4 (motor) and BA39 (extramotor; language). Microglia positive for FKBP5 or NF-*κ*B are indicated with blue arrows, other positive glia are indicated with orange arrows, and other negative glia are indicated with white arrows. (d) Increased magnification images of FKBP5+Iba1 and NF-*κ*B +Iba1 staining, with positive/negative glia indicated as described in b. (e) Nuclear/cytoplasmic NF-*κ*B intensity ratio quantification for neuronal (left), grey matter glial (middle) and white matter glial (right) staining between NPS1 and NPS2, stratified by brain region. * *p < 0*.*05*.

### BDNF expression correlates significantly with disease duration in C9-ALS

To explore relationships between expression of the 20 differentially expressed genes in C9-ALS and disease progression, gene expression was correlated with disease duration and ALSFRS slope of decline, identifying a positive correlation between *BDNF* expression and disease duration (Spearman’s *R = 0*.*64, p = 0*.*047*) (Figure 4a). BaseScope^™^ *in situ* hybridisation was used to validate our finding that *BDNF* expression correlates with disease duration in C9-ALS. *BDNF* expression in BA4 was manually graded using a product score to account for both cell and regional transcript abundance. *BDNF* was predominantly expressed in neurons, with heterogeneous abundance, at both the cell and regional level (Figure 4b). We confirmed a positive correlation between *BDNF* expression and disease duration (Figure 4c). Individuals with a short disease duration (i.e., less than 48 months post-onset)^54^ consistently showed lower levels of *BDNF* expression, while individuals with long disease duration (i.e., more than 48 months post-onset) exhibited higher expression.

**Figure 4.**
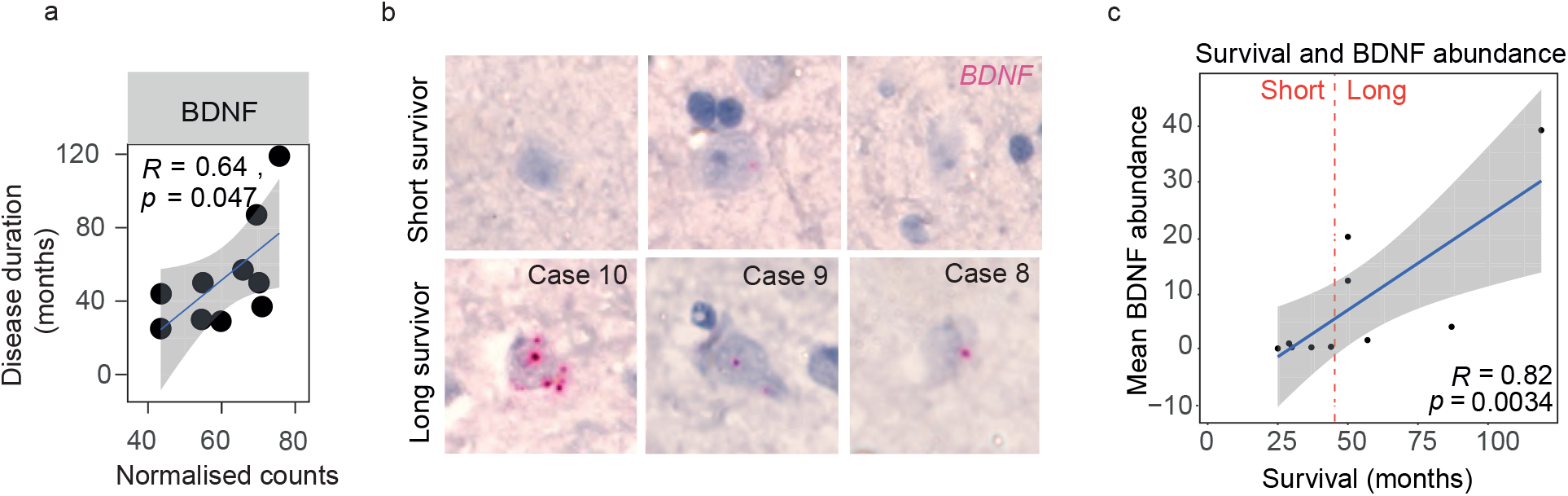
BDNF expression correlates with disease duration in C9-ALS. (a) Significant correlation between *BDNF* expression and disease duration. Spearman’s R and p-value is shown with a linear regression line and 95% confidence interval. (b) Example images of BaseScope™ *in situ* hybridisation of *BDNF* probes in short and long survivors. (c) Significant correlation between mean *BDNF* transcript abundance (product score) and survival (i.e., disease duration (months)). Spearman’s R and p-value is shown with a linear regression line and 95% confidence interval.

### Distinct inflammatory signatures exist across C9-ALS, sALS and SOD1-ALS cohorts

To interrogate whether the observed inflammatory signatures were specific to C9-ALS or common across multiple ALS cohorts, we next applied the nCounter Neuroinflammation Panel to sALS and SOD1-ALS cohorts (Table 3). Differentially expressed genes between each cohort and controls were largely different, with no overlap of genes passing the FDR threshold present in all three cohorts (Figure 5a-b). GO analysis of these genes revealed both distinct and shared significant terms across cohorts related to immune function and proteostasis, as well as other pathways (Supplementary Materials). Distinct terms included response to interleukin, microRNA gene transcription, neuronal death, post-translational protein modification, aggrephagy and chaperone-mediated protein transport in C9-ALS; cell development and morphogenesis in sALS; and translation initiation, protein kinase B signalling, chaperone-mediated protein folding in SOD1-ALS. Overlap included postsynaptic neurotransmission in C9-ALS and sALS, glial migration in C9-ALS and SOD1-ALS, and chemokine-mediated signalling, and T cell, B cell, and natural killer cell processes in sALS and SOD1-ALS (Supplementary Materials). Despite these differences, heatmap cluster analysis of C9-ALS, sALS, and SOD1-ALS cases using the filtered NPS gene list revealed two distinct NPS subgroups, present across the included cohorts, again delineated by the expression of two gene clusters related to immune response or axonal transport and synaptic processes (Figure 5c; Supplementary Figure 2a-b for full panel and GO). The two subgroups also did not appear to segregate clearly based on our available clinical metrics for cognitive function or glial pTDP-43 burden, unlike what was observed within the C9-ALS cohort alone. Interestingly, differential expression analysis revealed significant dysregulation of immune response genes (i.e., complement and microglial genes) in cognitively impaired cases (C9-ALS, sALS) (Supplementary Figure 2c-d) while only one significantly dysregulated gene, *CNN2*, was detected between unimpaired cases (C9-ALS, sALS) and controls. As such, it is possible that cognitively impaired cases have convergent disease mechanisms despite being from different cohorts, while unimpaired cases may be too diverse to detect significant dysregulation in this context.

**Table 3.**
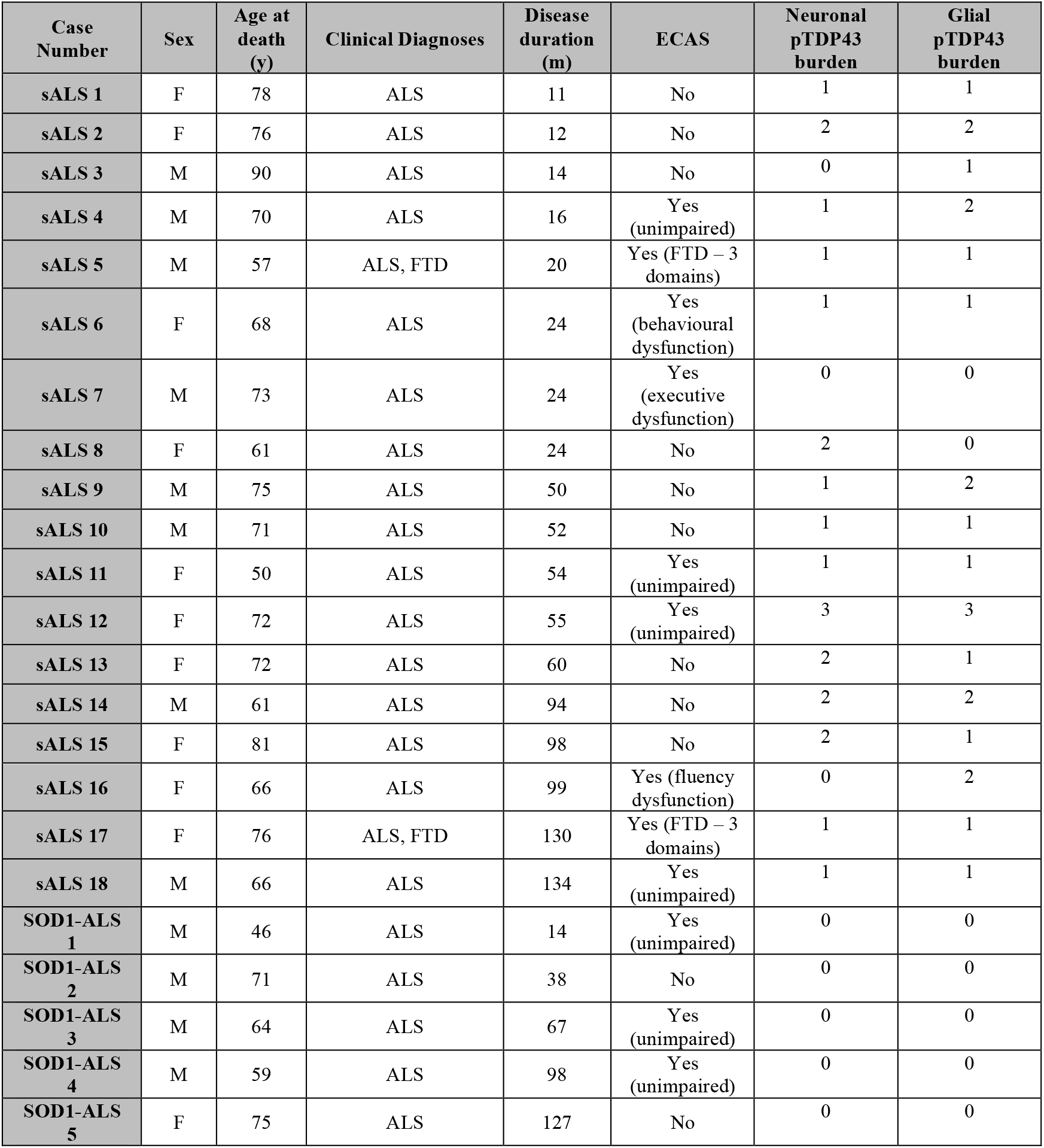
sALS and SOD1-ALS cohort demographics.

| Case Number | Sex | Age at death (y) | Clinical Diagnoses | Disease duration (m) | ECAS | Neuronal pTDP43 burden | Glial pTDP43 burden |
| --- | --- | --- | --- | --- | --- | --- | --- |
| sALS 1 | F | 78 | ALS | 11 | No | 1 | 1 |
| sALS 2 | F | 76 | ALS | 12 | No | 2 | 2 |
| sALS 3 | M | 90 | ALS | 14 | No | 0 | 1 |
| sALS 4 | M | 70 | ALS | 16 | Yes (unimpaired) | 1 | 2 |
| sALS 5 | M | 57 | ALS, FTD | 20 | Yes (FTD – 3 domains) | 1 | 1 |
| sALS 6 | F | 68 | ALS | 24 | Yes (behavioural dysfunction) | 1 | 1 |
| sALS 7 | M | 73 | ALS | 24 | Yes (executive dysfunction) | 0 | 0 |
| sALS 8 | F | 61 | ALS | 24 | No | 2 | 0 |
| sALS 9 | M | 75 | ALS | 50 | No | 1 | 2 |
| sALS 10 | M | 71 | ALS | 52 | No | 1 | 1 |
| sALS 11 | F | 50 | ALS | 54 | Yes (unimpaired) | 1 | 1 |
| sALS 12 | F | 72 | ALS | 55 | Yes (unimpaired) | 3 | 3 |
| sALS 13 | F | 72 | ALS | 60 | No | 2 | 1 |
| sALS 14 | M | 61 | ALS | 94 | No | 2 | 2 |
| sALS 15 | F | 81 | ALS | 98 | No | 2 | 1 |
| sALS 16 | F | 66 | ALS | 99 | Yes (fluency dysfunction) | 0 | 2 |
| sALS 17 | F | 76 | ALS, FTD | 130 | Yes (FTD – 3 domains) | 1 | 1 |
| sALS 18 | M | 66 | ALS | 134 | Yes (unimpaired) | 1 | 1 |
| SOD1-ALS 1 | M | 46 | ALS | 14 | Yes (unimpaired) | 0 | 0 |
| SOD1-ALS 2 | M | 71 | ALS | 38 | No | 0 | 0 |
| SOD1-ALS 3 | M | 64 | ALS | 67 | Yes (unimpaired) | 0 | 0 |
| SOD1-ALS 4 | M | 59 | ALS | 98 | Yes (unimpaired) | 0 | 0 |
| SOD1-ALS 5 | F | 75 | ALS | 127 | No | 0 | 0 |

**Figure 5.**
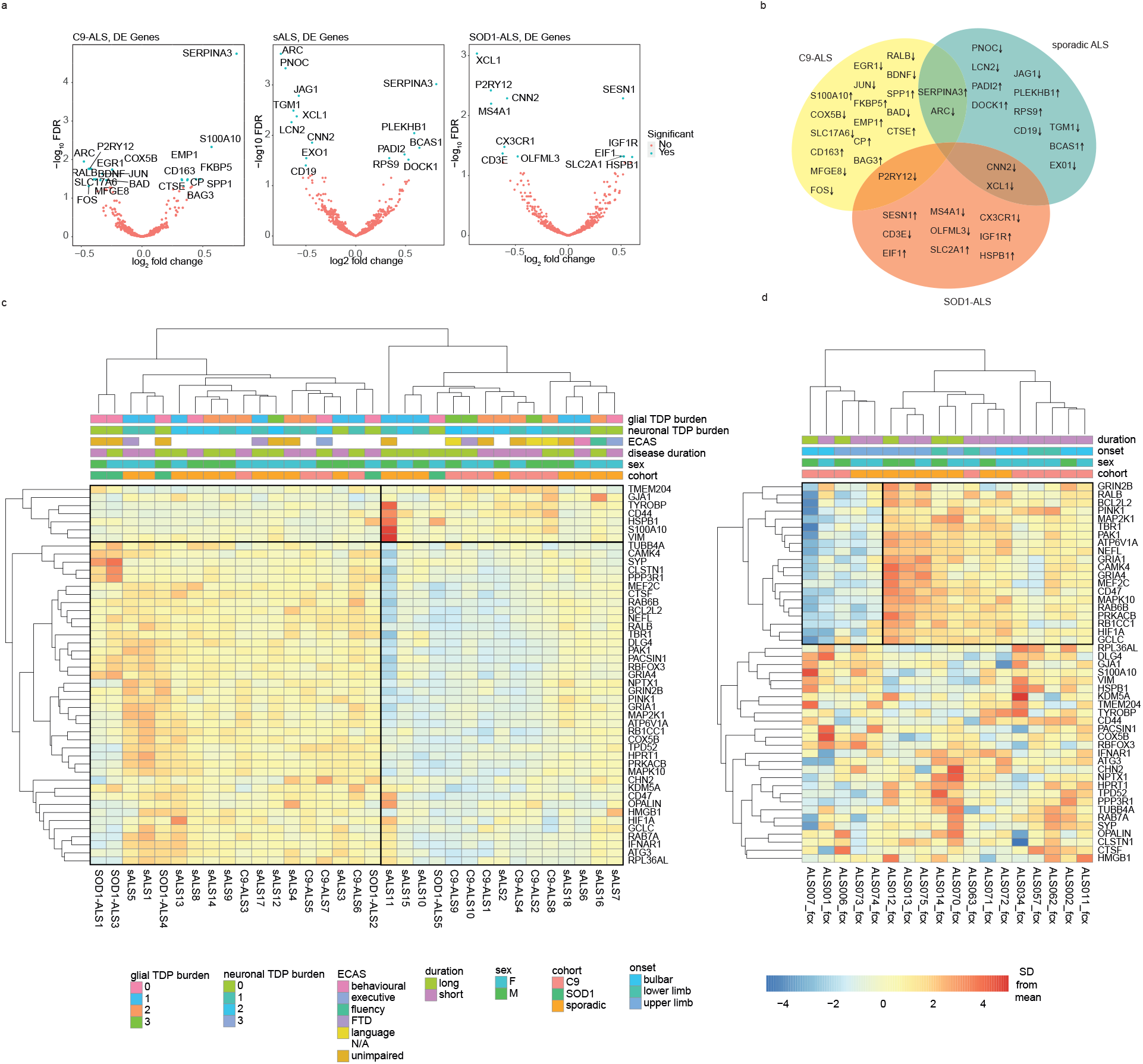
C9-ALS, sALS and SOD1-ALS cohorts exhibit both overlapping and unique neuroinflammatory signatures. (a) Volcano plot showing differentially expressed genes for C9-ALS (*n=10*), sALS (*n=18*) or SOD1-ALS (*n=5*) compared to controls by log2 fold change and -log10 p-value. Non-significant genes are represented in red and significant genes surpassing the Benjamini-Hochberg false discovery rate (FDR) threshold (i.e., p-adjusted < 0.05) are represented in blue. (b) Venn diagram showing the overlap of differentially expressed genes from (a) across cohorts, with arrow indicating direction of differential expression. (c) Clustered heatmap with filtered NPS gene list from Figure 1f, showing two distinct neuroinflammatory signatures across C9-ALS, sALS, and SOD1-ALS cohorts. Gene clusters 1 & 2 are boxed, with opposite directions of expression. Demographic (cohort, sex), clinical (disease duration, ECAS) and pathological (pTDP-43 burden) keys are shown. (d) Clustered heatmap with filtered NPS gene list from Figure 1f, showing two distinct neuroinflammatory signatures in the frontal cortex across C9-ALS and sALS cases from an independent, publicly available dataset, delineated particularly by the the 20 genes comprising the top quadrants of the heat map. Key is the same as in (c).

To test the generalisability of the filtered NPS gene list from C9-ALS cases across ALS cohorts, the list was next applied to the clustering analysis including cases from all cohorts (Figure 5a). Genes clustered in the same way as in the C9-ALS analysis (Figure 1e), and groups were delineated such that C9-ALS cases remained divided as previously. Opposite directions of expression were most apparent in the 40 genes encompassed within the clades comprising the lower quadrants of the heatmap (Figure 5c). The gene list was next tested further on an independent cohort of *post-mortem* frontal cortex and cerebellum of eight C9-ALS and ten sALS cases from publicly available RNA sequencing data^26^ (Figure 5d). Strikingly, in the frontal cortex, clustering analysis revealed a very distinct delineation between NPS1 and NPS2, present across both cohorts and particularly defined by the direction of expression of 20 of the genes in the filtered list (Figure 5c). Notably, this effect did not persist in the cerebellum, for which the clustering analysis using the filtered gene list did not reveal distinct subgroups (Supplementary Figure 2e), highlighting the possibility of region-specific inflammatory signatures.

## Discussion

This study investigated neuroinflammatory differences in deeply clinically phenotyped ALS *post-mortem* tissue, allowing us to compare molecular data directly with motor and cognitive as well as immunohistochemical features. We found an upregulation of microglia-specific genes in C9-ALS, substantiating previous findings of microglial dysregulation in cases with this genetic background^17,24^. Microglia have been shown to require C9orf72 for normal function in C9orf72 -/- microglia and peripheral myeloid cell models, demonstrating a pro-inflammatory response as a result of C9orf72 knockout^22,23^. Thus, haploinsufficiency resulting from *C9orf72* HRE may lead to the microglial dysregulation observed both here and elsewhere. In line with this, expression of many hits was shown to correlate with microglial and pTDP43 staining quantification.

Two genes whose expression was found to correlate with clinical scores (*BDNF, FKBP5*) were further validated with spatial resolution using immunohistochemistry or BaseScope^™^ *in situ* hybridisation. C9-ALS-related increased nuclear localisation, and thus activation of FKBP5 signalling partner, NF-κB, was observed in neurons and glia in both motor and extramotor regions, along with exhibition of microglia-specific NF-κB staining in white matter. Activation of NF-κB in glia is indicative of an upregulation of an inflammatory response mediated by IKK*α*/*β* kinases^55^, which are negatively regulated by autophagy^56,57^. It is possible that the more pro-inflammatory signature, increased BA4 grey matter glial NF-κB activation, and higher TDP-43 burden seen in NPS1 is related to lower levels of negative regulation via autophagy. Indeed, the *NLRP3* inflammasome, as well as several autophagy-related genes (i.e. *ATGs*), are part of Gene Cluster 1; activation of *NLRP3* is also increased with autophagy deficiency^57–59^. Thus, therapeutic studies involving the use of autophagy-targeting drugs may seek to consider stratification of cases based on inflammatory signatures to ensure the meaningful measurement of outcomes.

The utility of transcriptome data for identifying molecular signatures that correlate with survival has been recently demonstrated identifying a subgroup of ALS patients with poorer survival with differential expression of genes relating to oxidative phosphorylation^60,61^. Here we identify *BDNF* expression as a clinical correlate of survival, highlighting the additional mechanistic insights afforded through our targeted approach. Expression of *BDNF*, or brain-derived neurotrophic factor, in the motor cortex was found to be downregulated in disease, and positively correlated with disease duration, suggesting a protective effect. BDNF signalling has been previously demonstrated to have either neuroprotective^62^ or indirectly excitotoxic effects^63^. *BDNF* expression has also been shown to correlate with decreased cognition^64^. Importantly, many preclinical studies investigating the effects of BDNF in ALS are biased toward SOD1 mouse models^65,66^, in which increased BDNF-TrkB is observed^67^. Further, a phase III clinical trial conducted using recombinant methionyl human BDNF did not demonstrate therapeutic benefit, though this study and further trials conducted thereafter did not stratify genetically; importantly, *C9orf72* mutations in ALS had not been discovered at the time^68,69^. In contrast to SOD1 models, this study shows that *BDNF* expression is downregulated in C9-ALS *post-mortem* tissue, and C9-ALS cases appear to have a more inflammatory background; thus, BDNF-related treatments may function differently in a C9-ALS context. Expression of *BDNF* by immune cells was found to promote neuronal survival in human tissue culture^70^. Moreover, subcutaneous perfusions of BDNF have been shown to reverse microglial activation in aged mice^71^, perhaps through indirect downregulation of microglial MHC-II expression^72^. As such, cell type-specific manipulation of *BDNF* expression may provide a more nuanced approach to controlling microglial activation and neuronal loss. In addition to a possible treatment, BDNF could also have utility as a biomarker for disease prognosis. Recently, BDNF and pro-BDNF levels in CSF were shown to be associated with survival in ALS patients; in line with our discussion, C9-ALS patients showed significantly lower serum BDNF levels than non-carriers^73^.

While previous studies have used RNA sequencing to investigate and identify important gene expression signatures across the transcriptome^26,60^, our targeted approach to investigating neuroinflammatory signatures without amplification bias ensured a focused evaluation of neuroinflammation specifically. We identified two distinct molecular profiles, NPS1 and NPS2, with immune response terms enriched in Gene Cluster 1 and axon transport and synaptic signalling terms enriched in Gene Cluster 2. These signatures do not segregate clearly with known demographic, clinical, or pathological data in our cohorts suggesting that these signatures are not readily identifiable through visible features. These signatures are present in multiple ALS cohorts, within our study and in an independent publicly available dataset, underscoring their generalisability and, crucially, highlighting the importance of molecular stratification in clinical trials. Clinical trials may benefit from employing stratification methods based on molecular markers rather than, or in addition to, genetic and clinical criteria, as without stratification a positive effect of treatments on a particular subgroup may be obscured. For example, the recent macrophage-targeted sodium chlorite trial (NP001) showed no overall effect on the primary outcome measure^74^. However, subsequent subgroup analysis showed that those that did have a beneficial therapeutic response to the drug had higher than average levels of circulating IL-18 and LPS (akin to our NPS1), implying that molecular stratification by key circulating inflammatory markers could enable us to treat a subset of ALS patients for whom inflammation plays a more substantial role^74^. Our data would suggest that a combinatorial blood-based biomarker approach^75^, using circulating markers derived from a gene panel such as ours, validated across distinct ALS patient populations (as in Figure 5d), would be a more appropriate way to identify subgroups that would benefit from targeted therapies. Promising candidates are based on the 20 genes from our NPS-defining gene list that strongly delineate clusters in an independent, publicly available dataset (i.e., *GRIN2B, RALB, BCL2L2, PINK1, MAP2K1, TBR1, PAK1, ATP6V1A, NEFL, GRIA1, CAMK4, MEF2C, CD47, MAPK10, RAB6B, PRKACB, RB1CC1, HOF1A, GCLC*) and two additional NPS-defining genes (*CD44* and *TYROBP*) that also appear in a recently identified gene list defining a molecular subgroup relating to glial activation^61^. Molecular stratification, in the form of tissue derived and circulating biomarkers, is the mainstay of patient stratification for clinical trials in oncology^76^; given the convergence of these studies with our data, molecular subtyping should be considered for future trials implementing targeted therapies in people with ALS.

## Acknowledgments

This research was funded in part by the Wellcome Trust (108890/Z/15/Z) to OMR, a Pathological Society and Jean Shanks Foundation grant (JSPS CLSG 202002) to JMG and JOS, an NIH grant (5-R01-NS127186-02) to JMG, FMW, and JOS, a Motor Neuron Disease (MND) Scotland grant to JMG and CRS (2021/MNDS/RP/8440GREG), and a Sir Henry Dale Fellowship jointly funded by the Wellcome Trust and the Royal Society (215454/Z/19/Z) to CRS. For the purpose of open access, the author has applied a CC BY public copyright licence to any Author Accepted Manuscript version arising from this submission. This work would not be possible without the resources of the Edinburgh Brain Bank. The authors declare no conflicts of interest.

