## Supplementary Materials for "Distinct neuroinflammatory signatures exist across genetic and sporadic ALS cohorts"

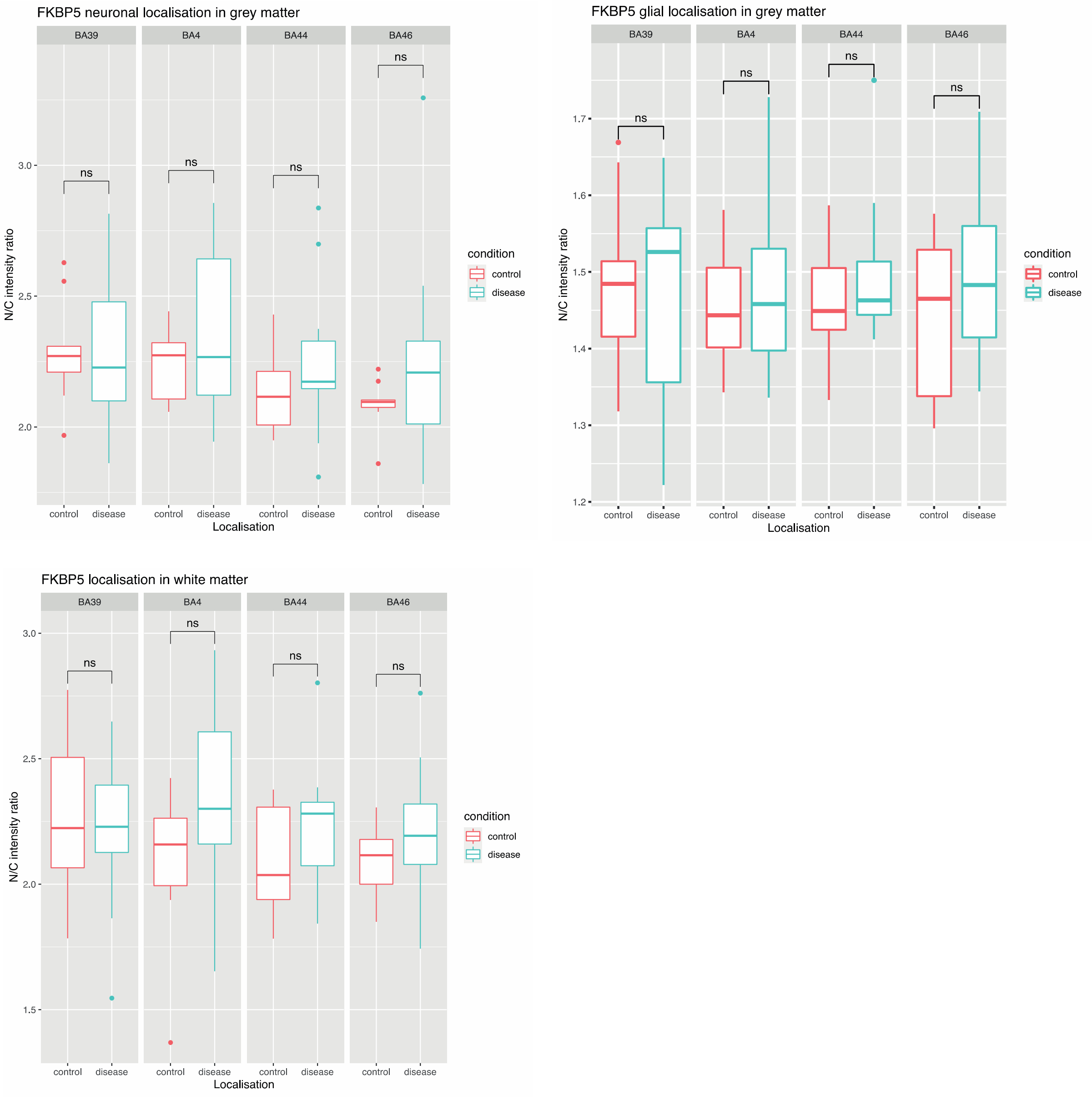


### Supplementary Figure 1. Supplementary FKBP5 analyses.

Nuclear/cytoplasmic FKBP5 intensity ratio quantification for neuronal, grey matter glial, and white matter glial staining between ALS and controls, stratified by brain region. * $p < 0.05$.

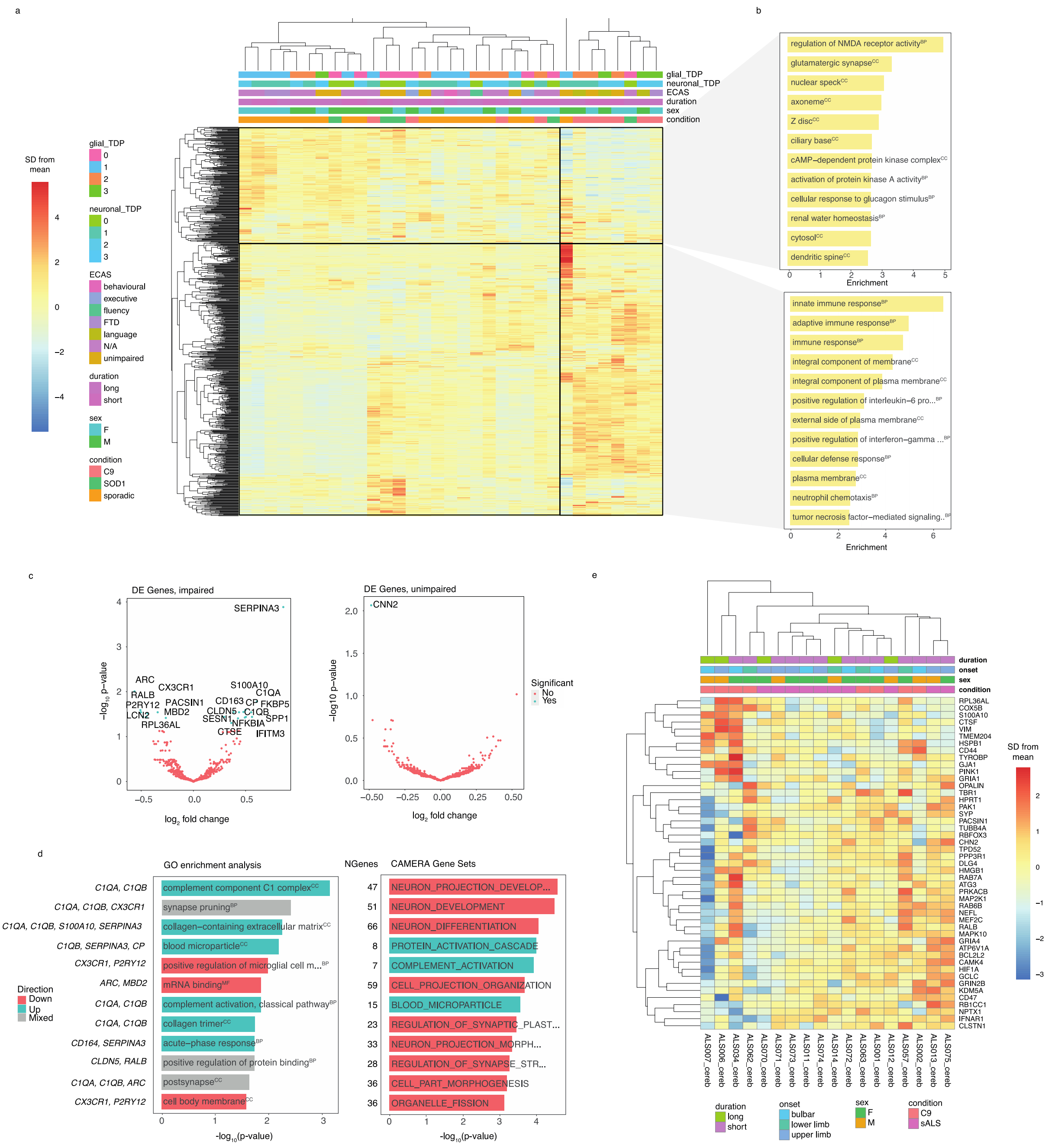


### Supplementary Figure 2. Supplementary heatmap analyses.

(a) Clustered heatmap with entire neuroinflammation panel (770 genes), showing two distinct neuroinflammatory signatures across C9-ALS, sALS, and SOD1-ALS cohorts. Demographic (cohort, sex), clinical (disease duration, ECAS) and pathological (pTDP-43 burden) keys are shown. (b) GO enrichment analysis for gene clusters that define signatures 1 and 2 showing top 12 dysregulated gene sets. Italicised terms are downregulated. (c) Volcano plot showing differentially expressed genes for cognitively impaired (C9-ALS and sALS) or cognitively unimpaired (C9-ALS and sALS) cases compared to controls by $\log_2$ fold change and $-\log_{10}$ FDR. (d) (left) GO enrichment analysis of genes enriched in cognitively impaired cases by type with $-\log_{10}$(p-value) score showing top 12 dysregulated gene sets; MF, molecular factor; CC, cellular component; BP, biological process. Italicised terms are downregulated; key genes for each term are shown to the left; (right) CAMERA gene set analysis of gene sets enriched in cognitively impaired ALS cases showing top 12 dysregulated gene sets, with the number of genes for each term shown to the left. (e) Clustered heatmap with filtered NPS gene list from Figure 1f, showing the expression of neuroinflammatory genes in the cerebellum across C9-ALS and sALS cases from an independent, publicly available dataset, delineated particularly by the first 20 genes.

**Online supplementary data on figshare.com**

Rifai, Olivia (2023): 1. Digital pathology analysis scripts. figshare. Online resource. https://doi.org/10.6084/m9.figshare.21916557.v1

Rifai, Olivia (2023): 2. Raw images from digital pathology analysis. figshare. Figure. https://doi.org/10.6084/m9.figshare.21916575.v2

Rifai, Olivia (2023): 3. Raw and normalised NanoString counts. figshare. Dataset. https://doi.org/10.6084/m9.figshare.21916581.v1

Rifai, Olivia (2023): 4. Differential expression analysis results. figshare. Dataset. https://doi.org/10.6084/m9.figshare.21916629.v1

Rifai, Olivia (2023): 5. GO enrichment and GSA results. figshare. Dataset. https://doi.org/10.6084/m9.figshare.21916656.v1

Rifai, Olivia (2023): 6. C9-ALS gene clusters list. figshare. Dataset. https://doi.org/10.6084/m9.figshare.21916671.v1
